# A point mutation in HIV-1 integrase redirects proviral integration into centromeric repeats

**DOI:** 10.1101/2021.01.12.426369

**Authors:** Shelby Winans, Stephen P. Goff

## Abstract

Retroviruses utilize the viral integrase (IN) protein to integrate a DNA copy of their genome into the host chromosomal DNA. HIV-1 integration sites are highly biased towards actively transcribed genes, likely mediated by binding of the IN protein to specific host factors, particularly LEDGF, located at these gene regions. We here report a dramatic redirection of integration site distribution induced by a single point mutation in HIV-1 IN. Viruses carrying the K258R IN mutation exhibit more than a 25-fold increase in integrations into centromeric alpha satellite repeat sequences, as assessed by both deep sequencing and qPCR assays. Immunoprecipitation studies identified host factors that uniquely bind to the mutant IN protein and thus may account for the novel bias for integration into centromeres. Centromeric integration events are known to be enriched in the latent reservoir of infected memory T cells, as well as in patients who control viral replication without intervention (so-called elite controllers). The K258R point mutation in HIV-1 IN reported in this study has also been found in databases of latent proviruses found in patients. The altered integration site preference induced by this mutation has uncovered a hidden feature of the establishment of viral latency and control of viral replication.

## Main

Insertion of the viral DNA genome into the host cell genome, a process termed integration, is an obligate step of a successful retroviral infection. By permanently integrating the viral genome into the host genome, retroviruses are able to persist indefinitely in the infected cell as a provirus. Integration is solely catalyzed by the virally encoded integrase (IN) protein^1,2^. Although all of the host genome is available as a target for integration at some frequency, the distribution of integration sites across the genome is not completely random^3–5^, and various retroviruses exhibit distinct integration site preferences^6^. Specifically, human immunodeficiency virus (HIV-1) has a preference for integrating into active gene regions^7^. Differential integration site selectivity can be primarily explained by the binding of the viral IN protein to various host factors^8^. The preference for HIV-1 to integrate into active genes for instance was found to be in part due to binding of the IN protein to the host factor LEDGF, a general transcriptional activator^9–12^. These host factors are believed to act largely as bimodal tethers, binding both the viral IN protein and host chromatin, and thereby biasing integration sites to specific genomic regions^13^.

Integration targeting by chromatin tethering is a conserved mechanism amongst retroviruses and retrotransposons alike. The yeast Ty elements in particular exhibit highly specific integration targeting, down to the nucleotide in some cases14. Ty5 elements are mainly integrated into heterochromatic regions such as telomeres or the mating type loci through interaction of the Ty5 IN protein with the yeast silencing factor Sir4^15,16^. The affinity of Ty5 IN for Sir4 is dependent on phosphorylation of the targeting domain of IN^17^. In the absence of IN phosphorylation, as occurs during certain stress conditions, Sir4 binding is lost and Ty5 integration is dramatically redirected in a dispersed fashion throughout the yeast genome^17^.

HIV-1 IN is known to be heavily post-translationally modified, but no evidence to date has linked any post-translational modifications (PTMs) to integration site selection18,19. There are four major acetylation sites in the C-terminal domain (CTD) of HIV-1 IN (K258, K264, K266 and K273)20,21. We mutated these lysine residues to charge-conservative arginines, either singly or in combination. We generated pseudotyped single-round infection HIV-1 viral reporter constructs expressing luciferase, packaged into virion particles with either a WT IN or a mutant IN, and used them to transduce cells in culture. Infected cells were collected at 48 hours post-infection and assayed for successful viral transduction by quantifying viral DNA products as well as luciferase activity (Fig. 1). We have previously reported the effects of these mutations on viral transduction, and that mutation of all acetylated lysine residues in combination led to a dramatic decrease in proviral transcription immediately after viral DNA integration^22^. In this study, we focus specifically on the K258R point mutation in HIV-1 IN.

**Figure 1:**
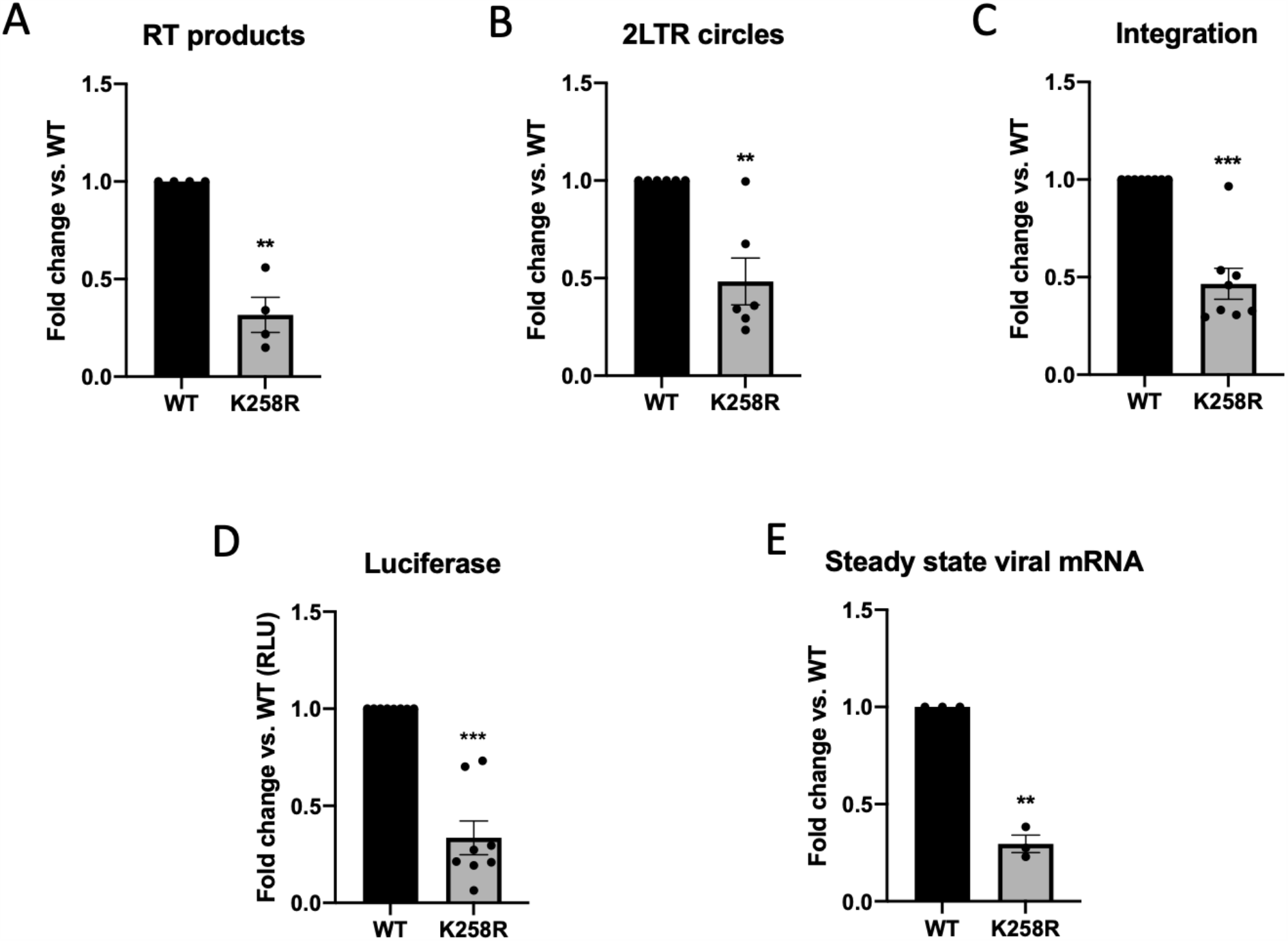
K258R point mutation in HIV-1 IN has modest effects on early viral replication. HeLa cells were infected with virus generated from pNL4.3.Luc.R-E-carrying either WT or K258R mutant IN. Infected cells were collected at 2 days post-infection. Abundance of (A) reverse transcription (RT) products and (B) 2-LTR circles was determined by qPCR and normalized to a housekeeping gene (n=4 and 6 respectively). (C) Proviral integration frequency was assayed using a nested PCR Alu-gag approach (n=7). (D) Luciferase activity was measured (RLU) and normalized by protein content to adjust for number of cells in input sample (n=8). (E) Steady state viral mRNA levels were measured by qPCR of infected cellular cDNA using primers against spliced *tat* message (n=3). All data is shown as a fold change relative to WT and is the average of the indicated number of independent biological replicates +/- SEs. Statistical significance was gauged by two-tailed paired t-test (*p<0.05, **p<0.01, ***p<0.001).

The K258R point mutation in IN caused a modest 3-fold defect in total reverse transcription (RT) as gauged by qPCR quantification of viral DNAs (Fig. 1A), and an equally modest 2-fold decrease in the abundance of 2-LTR circular DNA, a structure generated upon nuclear entry (Fig. 1B). The mutation resulted in a similar 2-fold reduction in the levels of proviral DNA formed after infection as compared to WT, measured by qPCR amplification of host-viral junctions (so-called Alu-gag assays; Fig. 1C). Quantification of luciferase activity and steady state viral mRNA transcripts corroborated a modest decrease in overall viral transduction (Fig. 1D-E). These findings indicate that all viral DNA intermediates and viral mRNA levels are reduced by a comparable amount in the cells infected with virus carrying the K258R IN mutation, and that there is no significant defect at the step of integration. The small decrease in transduction is accounted for by the initial decrease in reverse transcription products and thus in viral DNA available for subsequent steps.

Based on the alteration of integration site distribution induced by changes in phosphorylation status of the retrotransposon Ty5 IN in yeast, we mapped integration sites produced by the acetylation mutant INs as compared to WT IN. We used PCR and high-throughput DNA sequencing methods to recover and characterize viral-host genome junctions. Integrations were then mapped to unique human sequences using Bowtie2 and analyzed for correlation with RefSeq genes, CpG islands, transcription start sites, DNase hypersensitivity sites and various protein or histone binding sites identified in ChIP-seq datasets. These alignments are restricted to single- or low-copy number genomic sequence databases.

The combinatorial quadruple acetylation (QA) mutant IN and three of the four point mutant INs produced proviral integration patterns with very little deviation from WT pattern (Fig. S1). However, we observed significant differences in the distribution of proviruses integrated at uniquely mapped sequences by the K258R mutant IN as compared to those formed by WT IN (Fig. 2, Table 1). As previously shown, WT HIV-1 proviruses were preferentially located in and around annotated RefSeq genes. The K258R mutation reduced this preference for integration into genes to the level of random chance (matched random control, MRC) (Fig. 2A). Similarly, the WT IN showed the expected slight preference for integrating near CpG islands, but the K258R mutant IN showed less of this preferential targeting (Fig. 2B). This general reduction in integration frequency near these sites held true for other genomic features as well, including DNase hypersensitivity regions and RNA polymerase II binding sites (Fig. 2C-D). These decreases were not due to an overall decrease in integration frequency, since all quantifications were normalized to the total number of unique integrations mapped. The distribution of integration sites relative to transcription start sites, however, was unchanged by the K258R mutation (Fig. 2E). We also correlated proviral integration sites to the genomic coordinates of various pre-infection histone modifications present in HeLa cells (Fig. 2F). We observed no notable differences in the frequency of proviral integration sites occurring in proximity to any of four chromatin modifications (H3K27ac, H3K36me3, H3K4me3 and H3K9me3) generated by the K258R mutant IN as compared to WT (Fig. 2F).

**Figure 2.**
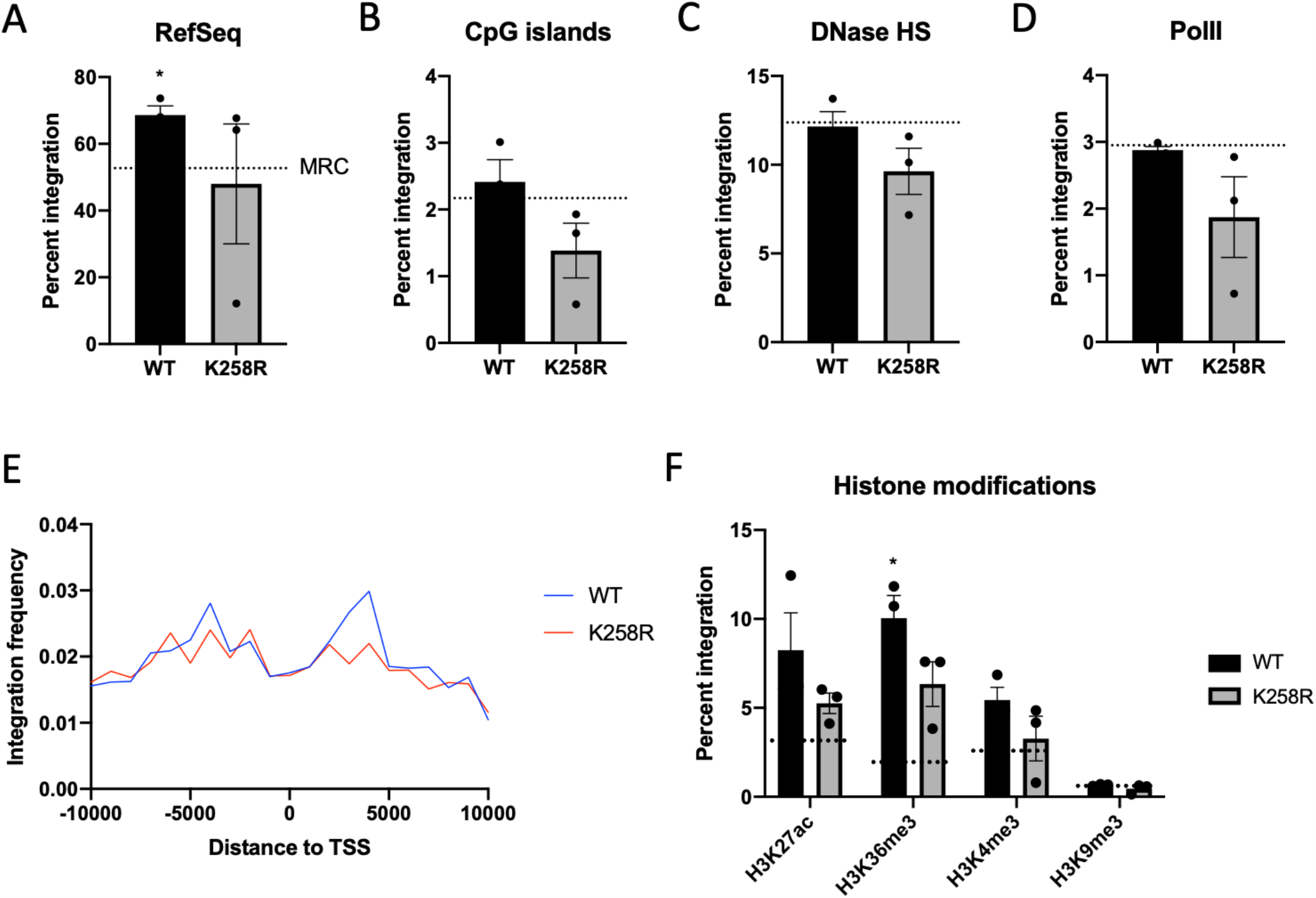
The K258R mutation in IN alters integration site distribution. Integration sites were mapped to the GRCh38 human reference genome assembly using Bowtie end-to-end alignment. Frequency of integrations falling within 1 kb of (A) RefSeq genes, (B) CpG islands, (C) DNase hypersensitivity sites and (D) RNA polymerase II binding sites was calculated using BedTools. The frequency of integrations expected to be located near these features by random chance (matched random control, MRC) is shown as a dashed line. (E) Distribution of integrations around transcription start sites (TSS). Integrations in a 10 kb window around TSS are shown. (F) Frequency of integrations within 1 kb of select pre-infection histone modification sites. Data shown is the average of three independent biological replicates +/- SEs. Statistical significance relative to MRC as gauged by a one-sample t-test is shown (* p<0.05). Additional statistical analysis comparing the integration site pattern of WT and K258R mutant IN is shown in Table S4.

**Table 1:** Number of unique integrations mapped (N=3 biological replicates)

|  | WT | K258R |
| --- | --- | --- |
| Unique integrations | 17636 | 18850 |
| Within RefSeq genes | 12363 | 11737 |
| TSS (5 kb) | 3630 | 3412 |
| TSS (1kb) | 671 | 387 |
| CpG islands (5 kb) | 2549 | 2291 |
| CpG islands (1 kb) | 426 | 316 |
| RNA Pol II (1 kb) | 501 | 447 |
| DNase HS (1 kb) | 2048 | 2018 |
| H3K27ac (1 kb) | 1197 | 1080 |
| H3K36me3 (1 kb) | 1656 | 1379 |
| H3K4me3 (1 kb) | 875 | 809 |
| H3K9me3 (1 kb) | 121 | 108 |
| Within centromeres | 146 | 1576 |

The analysis of the distribution of integrations of mutant K258R IN into unique mappable sites showed a loss of selective targeting to active genes as well as other features, but did not reveal a concomitant increase in integration frequency elsewhere. To examine the distribution of integration sites more globally and determine where the K258R mutant IN is being redirected, we made use of scan statistics to identify regions of the genome with high numbers of viral integrations in an unbiased fashion, and specifically including highly repetitive sequences ^23^. We analyzed common sites of integration or “hot-spots” using the custom perl script”^24,25^. This script first removes identical reads resulting from potential PCR duplication. Reads with identical viral-host genome junction sequences but disparate read lengths (breakpoints) were condensed into a single event. To account for potential copying errors induced by multiple rounds of PCR or sequencing we also combined those integrations in which the host sequence had >95% similarity over the length of the read. We then used a sliding window to scan the human genome for common sites of integration. For our purposes hot-spots were defined as 5 or more integrations in a 10 kb window.

We identified an unprecedented number of hot-spot sites for integration by the K258R mutant IN that all clustered in centromeres (Table 2). The frequency of insertion of the mutant into centromeric regions was extraordinarily high, with 10 clear genomic hot spots of integration. There were no such detectable integration hot-spots in cells infected with WT HIV-1 virus. WT HIV-1 IN has been previously reported to disfavor integration into centromeric repeats with on average less than 1% of detectable proviruses found in or near centromeres^7,26^. The clustering we observe in the K258R mutant integration distribution could not be attributed to selective outgrowth of the infected cells in the population as the samples were collected only 48 hours post-infection.

**Table 2:** Hot-spots of integration for viruses carrying the K258R IN mutation (5+ integrations in a 10 kb window)

| Genomic coordinates | Number of integrants |
| --- | --- |
| Chr14: 17749223-17757726 | 5 |
| Chr13: 17630007-17635451 | 6 |
| Chr21: 12443946-12448362 | 6 |
| Chr21: 12534522-12533844 | 6 |
| Chr13: 17669962-17678635 | 7 |
| Chr14: 18151557-18159208 | 7 |
| Chr22: 14673032-14672354 | 8 |
| Chr1: 125173987-125183192 | 9 |
| Chr22: 15024034-15019610 | 9 |
| Chr1: 143246863-143248277 | 12 |

To better quantify all integration events in centromeric regions, we extracted genomic coordinates of centromeres from the hg38 human reference genome and determined the distance from each integration to the nearest centromere. In agreement with the hot-spot analysis, we found a dramatic increase in integration frequency in centromeric regions specifically for proviruses integrated by the K258R mutant IN as compared to WT (Fig. 3A). We found that an average of 28% of the proviruses integrated by the K258R mutant IN occurred into centromeres. Again, we detect less than 1% of the proviruses integrated by a WT IN in centromeres, below even what is expected by random chance. The observed preference of K258R is specific for centromeric sequences, and we did not observe an increase in integration in the flanking peri-centromeric region (Fig. 3B).

**Figure 3:**
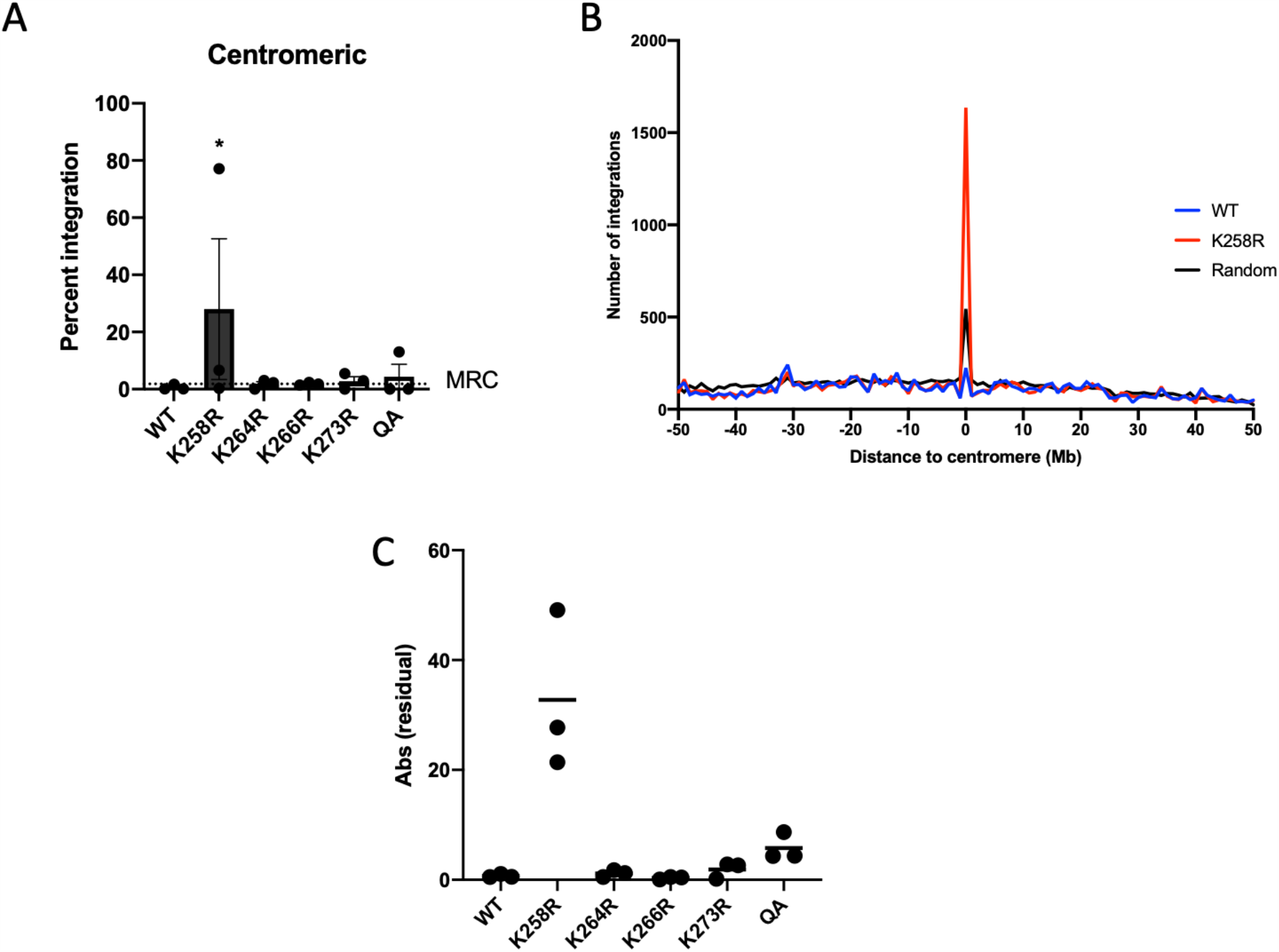
K258R mutant HIV-1 IN biases integration toward centromeres. (A) Number of integrations located in centromeric regions were normalized to total detected integration sites and are shown as a percent of the total. Integration frequency into centromeres in the matched random control (MRC) data set is shown as a dashed line. Data is shown as the average of three independent biological replicates +/- SEs. Statistical significance relative to MRC was calculated by one-way ANOVA corrected for multiple comparisons. (B) The distance to the nearest centromere was calculated for all WT and K258R mutant integration sites. A 50 Mb window flanking each centromere was segmented into 100 equal sized bins of 1 Mb. The number of integrations falling in each bin was quantified and is shown as a count (WT in black, K258R in red). (C) To assess variability of the altered integration centromere targeting phenotype we plotted the absolute residual from the mean for each independent trial. Statistical significance of variance was calculated using Levene’s test (*** p<0.0001, n=3).

We also analyzed the integration sites generated from the other acetylation mutant IN proteins. On average 1.7 – 4% of the detected proviruses integrated by these mutant IN proteins were detected in centromeric regions, a 2-4 fold increase as compared to WT (Fig. 3A). Thus, while all mutants exhibited a slight increase in preferential targeting to the centromeres, the K258R mutation alone strongly retargeted integration into centromeres at a shockingly high frequency, indicating that the effect of the K258R mutation in IN is unique to this residue, and not a general feature of blocking IN acetylation.

It should be noted that the magnitude of the observed phenotype in the NGS analysis was highly variable between independent replicate experiments. The fraction of the total integrations mediated by the K258R mutant IN recovered in centromeric regions ranged from extraordinarily high (∼80% -- the vast majority of integrations) to only moderately high (6% and 1%), but the proportion was consistently much higher than seen with the WT IN. To document this variability, we plotted the absolute value of the residuals from the mean observed in each replicate sequencing run for WT as well as in all IN mutants (Fig. 3C). The K258R mutation in IN produces a broad range of centromeric integration frequencies whereas WT IN and other IN mutant viruses gave a tight, uniform distribution around the mean in all trials. The potential for dramatically increased centromeric integration is a unique attribute of the K258R mutant IN.

The large variability of the observed integration targeting phenotype is likely attributable to how repetitive DNAs are sequenced and/or mapped. Traditional bioinformatics tools to map sequence data to the genome are limited in their capacity to deal with repetitive sequences, and many repeat elements are not even present in genome assemblies because they cannot be accurately placed. For this reason many integration site mapping studies to date focus exclusively on uniquely mapped reads to avoid the complexities of handling reads that map to multiple sites (“multi-mapping reads”). Our sequencing reads were mapped using a stringent Bowtie2 end-to-end alignment algorithm, with conservative reporting options that likely underestimate the true frequency of utilization of repetitive DNA as targets for integration. To obtain independent confirmation of the striking retargeting, we made use of several other bioinformatics tools commonly used in the field to re-analyze our integration site sequencing data.

We first confirmed this preference of the K258R mutant IN for integrating into centromeres using a Bowtie2-based sensitive local alignment strategy which allows for “soft-clipping” or omission of characters from the ends of reads in order to achieve the best alignment score. This can be advantageous if adaptor and/or viral sequences were not fully removed from the ends of reads in initial analysis steps but is in general a less conservative mapping approach. We further validated the centromeric integration preference of the K258R mutant IN using the BLAT mapping algorithm^27^, which is more commonly used amongst published integration site analysis studies. The BLAT mapping algorithm is based on BLAST and similarly reports all valid alignments above a set threshold score regardless of whether a read is unique or multi-mapping to repetitive sequences. Regardless of mapping algorithm, the data show that the K258R mutation in IN results in a dramatic redirection of integrations towards centromeres (Fig. S2). This site bias is not seen in any of the replicate tests of WT IN or other acetylation mutant IN proteins.

While the initial integration site mapping indicated that the K258R IN mutation induces a preference for integrating into centromeric regions, these algorithms do not identify specific target sequences and in fact do not even consider integration into the vast majority of repetitive sequences, which are largely excluded from the human reference genome. Centromeres are composed of tandem repeats, including both very short unit length repeats, and a high proportion of so-called alpha satellite sequence DNA comprised of alphoid repeats with a unit length of approximately 171 bp^28^. A number of other satellite sequences are present at lower abundance in the centromeric regions as well^26,29,30^. To determine which class of repeats may be specifically targeted by the K258R mutant IN, sequencing reads were mapped directly to the RepeatMasker track from the UCSC Genome Browser^31^. The RepeatMasker track includes all known repetitive sequences present in the genome, including simple repeats and shorter repeat units that are not present in the reference genome assembly. This allowed us to quantify integrations into all known repetitive regions, both in the centromere and outside, as well as obtain information on the repeat classes that are preferred targets of integration.

The K258R IN mutation causes a specific targeting of integrations into alphoid repeat sequences (Fig. 4A). This preference for alphoid repeats is not seen with WT IN or other acetylation mutant IN proteins, and indeed alpha satellite DNA specifically has been previously reported to be a disfavored target of WT HIV-1 integration^26^. The frequency of integrations into other common repeat classes such as Alu and L1 elements were not significantly different between WT and mutant INs (Fig. 4B). The K258R mutation of IN seems to uniquely redirect integrations to alpha satellite repetitive DNA and not other classes of repeat sequences.

**Figure 4:**
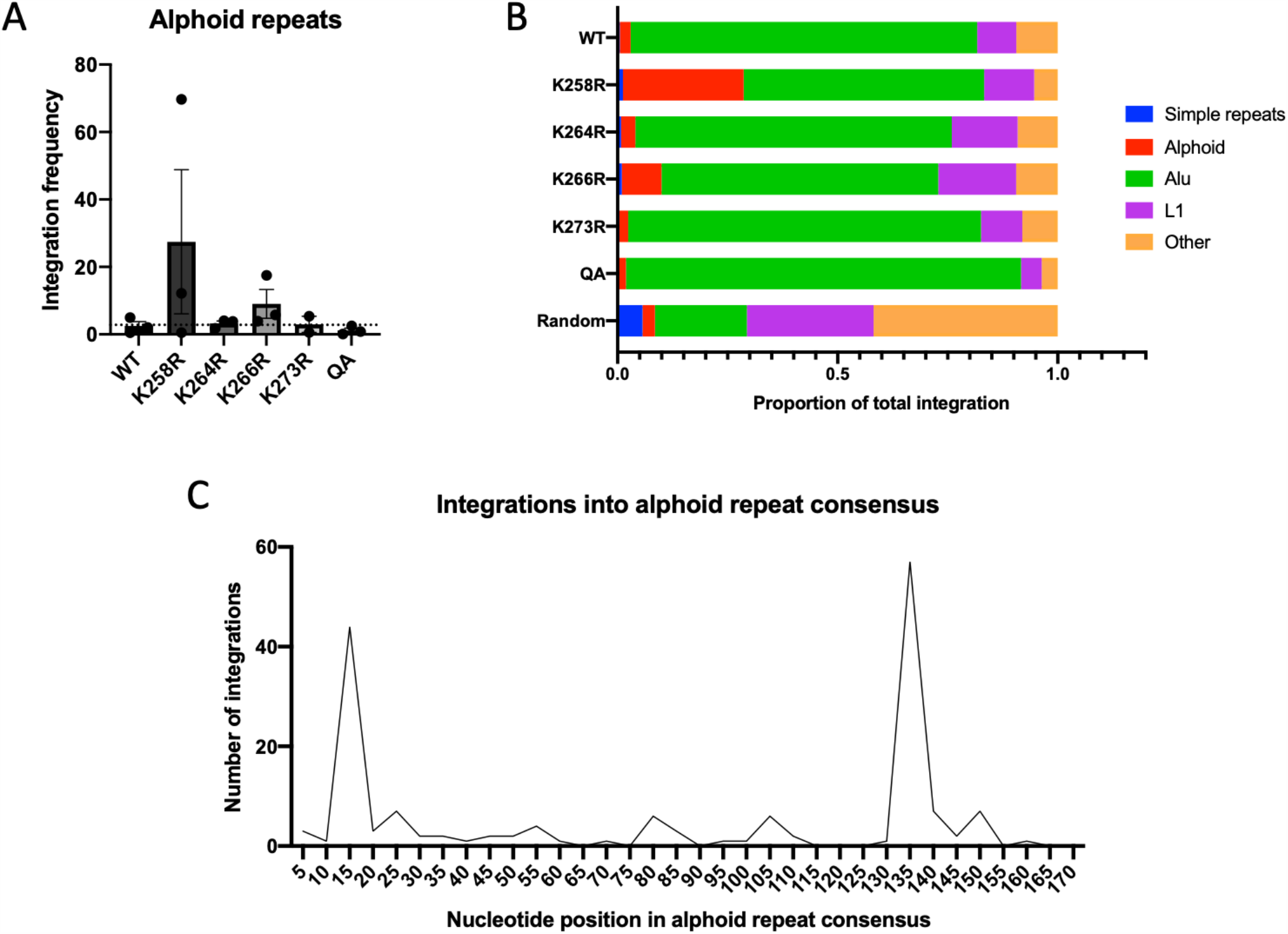
Mapping of proviral integrations to repetitive regions in the human genome. NGS reads from three independent biological replicate libraries were aligned to the RepeatMasker track from the UCSC genome browser. (A) The number of integrations mapping to alphoid DNA repeats was determined and normalized to the total number of mapped integrations and is shown as a percent of the total (n=3). The frequency with which integrations would be expected to fall in alphoid repeats if integration were random is shown as a dashed line (MRC). (B) The proportion of integrations that mapped to specific repeat elements relative to the total number of reads that mapped to the RepeatMasker track is shown. Only the most commonly targeted repeat elements are displayed. (C) Schematic of integration sites along the length of a single alphoid repeat. Unique host sequences immediately flanking each integration by the K258R mutant IN were aligned to an alphoid repeat consensus sequence (AJ131208.1) using Clustal Omega multiple sequence alignment. The consensus sequence was split into bins of 10 nucleotides and the number of integrations in each bin were counted. Shown are the integration counts falling in each bin summed over three replicates.

For all integrations by the K258R mutant IN that mapped to the centromere, we extracted the immediate flanking host genome sequences (10 bp upstream and 10 bp downstream), removed all identical junctions to be conservative and then aligned these to the alphoid repeat consensus sequence (AJ131208.1). We observe highly selective sites of insertion within the alpha satellite sequence by the K258R mutant IN protein, with two preferred spots of integration at nucleotide position 13 and 133 in the alphoid consensus sequence (Fig. 4C). The best alignment for the viral-host junction sequences at each hot spot was identical to the base position, but notably the host sequences at all the junctions were distinct, and thus represented distinct members of the alphoid repeat family. Thus, the many insertions into the alpha repeats are truly independent integration events. These two preferred sites in the repeat do not share a high level of sequence identity. There are no known protein binding motifs near either of the preferred sites. It is thus unclear why either of these two sequences is a preferred hot-spot for the mutant IN.

Due to the variability in the magnitude of the phenotype as well as the limitations of deep sequencing and available analytic tools, we wanted to verify the observed altered integration site distribution using a second method. In a modification of the Alu-gag method to quantify integration frequency, we devised a nested PCR approach to specifically assay for integrants in centromeric repeats. We replaced the primer located in the Alu repeat element that is typically used in Alu-gag assays with primers complementary to the alphoid repeat consensus sequence ^26,32^. We utilized two unique alphoid primers in our assay. To analyze both the 5′ and 3′ ends of the provirus we used primers complementary to either gag or luciferase respectively. This allowed for four unique combinations of primers in the first round of PCR that would selectively amplify proviruses in or near centromeric alphoid repeats. A subsequent second round quantitative PCR, using LTR specific primers, reported the yield of amplified viral DNA. The assays revealed a dramatic increase in the frequency of centromeric integration events for proviruses integrated by the K258R mutant IN (Fig. 5A). The magnitude of the effect was again highly variable, both between primer combinations and within a given primer pair, but was always dramatic. The K258R mutant IN increased integration frequency near centromeric alphoid repeats over the wild-type control by an average of 30-400 fold. The alpha satellite bias was again only seen with the K258R mutant. All other mutations blocking other acetylation sites displayed a similar level of centromeric integrations as wild type controls.

**Figure 5:**
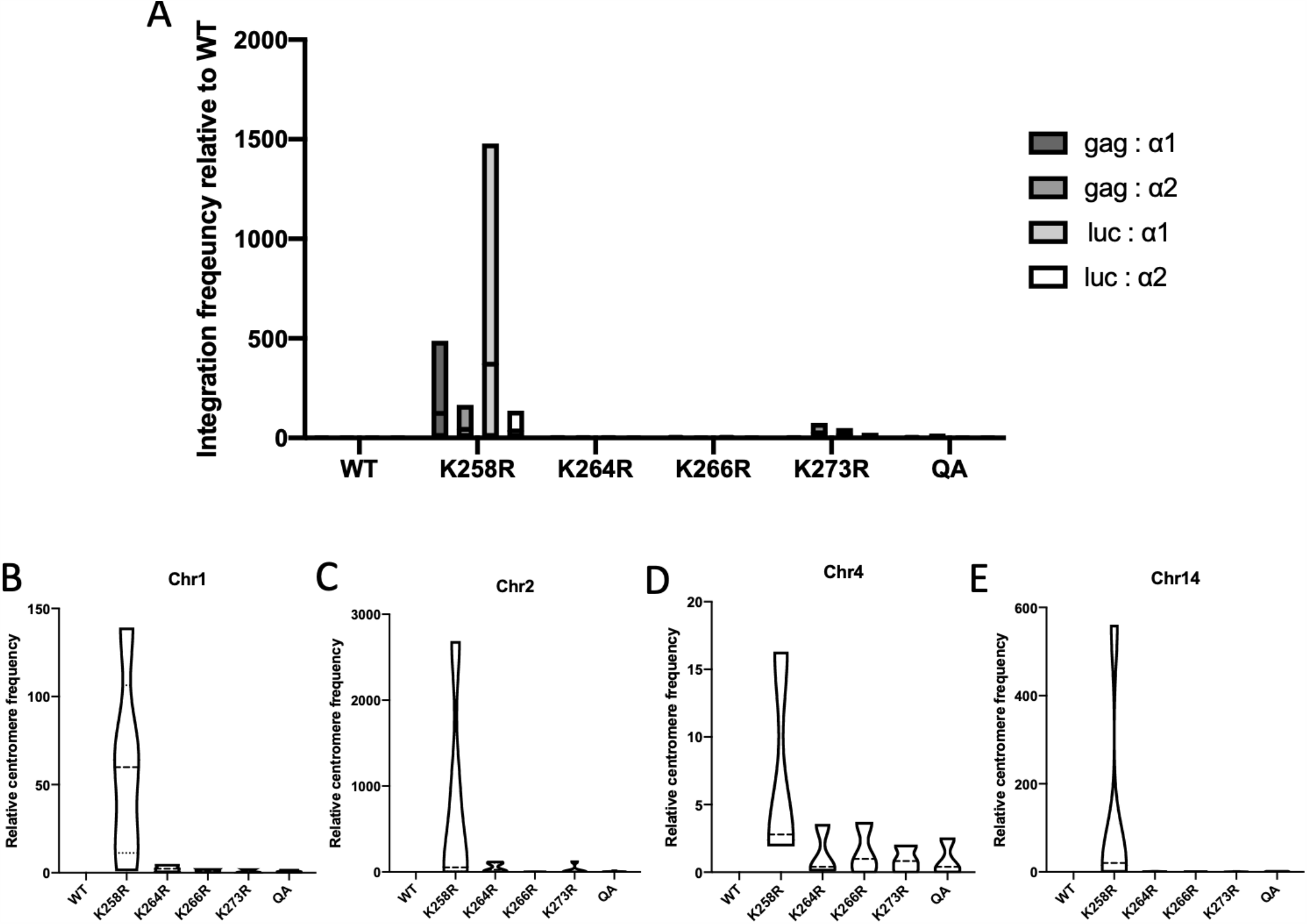
Quantification of integration frequency into centromeric regions by qPCR methods. (A) Integration into centromeric alphoid repeat DNA was quantified using a modified Alu-gag based nested PCR approach. Two unique primers were designed complementary to an alphoid repeat consensus sequence (α1, α2) and used instead of the typical Alu primer. Two primers at either end of the viral genome were used – either in the 5′ end of gag or in the 3′ UTR of the luciferase (luc) reporter gene. First nest PCR was performed with these four primer combinations. Shown are the results of a second nest quantitative PCR using LTR specific primers normalized to total integrated provirus levels as measured by Alu-gag PCR. Data from a minimum of three independent replicates is shown relative to WT as box plots to show the minimum, maximum and mean values. (B-E) Quantitative PCR using chromosome specific centromere primers. Viral LTR-host genome junctions were amplified and centromere content was subsequently quantified using qPCR with chromosome specific primers (see Table S1 for all primer sequences) and normalized to total integrated provirus levels as measured by Alu-gag PCR. Shown is the relative centromere content for each infected sample relative to WT from a minimum of three independent replicates presented as a violin plot to accurately represent the data distribution.

In our initial analysis to identify common sites of integration from NGS data, the identified genomic hot spots were all found in only a subset of chromosomes (Table 2). To determine whether the K258R mutant IN displayed any particular chromosomal preference, we also performed a qPCR assay utilizing chromosome-specific non-repetitive centromere primers to quantify specific centromeric DNA content present at the LTR-host genome junction. Shown are some representative examples using chromosome-specific primers for chromosomes 1, 2, 4 and 14 (Fig. 5B-E). The K258R mutant virus was observed to integrate much more frequently than WT or any other acetylation IN mutant viruses at sites near the centromeres regardless of chromosome. The apparent bias for some chromosomes that we observed in the initial “hot spot” analysis of the NGS data could be due to gaps or discrepancies in the assignments of the centromere sequences present in the genome assembly database. The PCR data suggest that K258R virus is targeted to centromeres of many, if not all, chromosomes.

Because HIV-1 integration is in part targeted through host factor interactions, it is plausible that the K258R mutation in the IN protein could modulate integration site selection by mediating differential binding of a specific host factor. To test this possibility, we generated mammalian expression vectors expressing either WT or K258R mutant IN protein and tested for host binding proteins. In both cases, the IN protein was N-terminally tagged with HA for immunoprecipitation. HEK293T cells were transfected with the IN plasmids and lysates were harvested after 24 hours of expression. Adequate and comparable expression of both WT and mutant IN proteins was confirmed via Western blot using both HA- and IN-specific antibodies. WT and mutant IN were immunoprecipitated and interacting host proteins were subject to mass spectrometry for identification.

We identified 43 and 56 proteins that bound to WT or K258R IN proteins, respectively, above the background of an empty vector control (Table 3). The majority of these host factors were shared between WT and K258R mutant IN. Based on a preliminary gene ontology analysis, the majority of factors that bind either WT or mutant IN protein are generic nucleic acid binding proteins (Table 3). Approximately a third of the 43 proteins we detected binding to WT IN have been previously reported by another mass spectrometry screen done in HEK293T cells, validating our approach^33^. We did not detect LEDGF in our immunoprecipitation, in agreement with previous studies that also failed to recover LEDGF in similar experiments in HEK293T cells^33^.

**Table 3:** Host proteins immunoprecipitated with WT or K258R mutant IN protein

| Sample(s) | Host protein names |
| --- | --- |
| WT/K258R | PRKDC, MDN1, MYBBP1A, NUP205, CAND2, GCN1, NUP188, CKMT1A, IMMT, HEATR1, IPO4, UBE3C, AIFM1, FANCI, ABCD3, ATP2A2, ABCE1, LTN1, SUCLA2, COQ8B, ATAD3C, DDX20, AFG3L2, ATAD3A, ATAD3B, RCN2, SGPL1, TYK2, SLC16A1, MCM7, TIMM50, ARF4, RRP12, PPP1CB, SLC25A10 |
| WT | TEX10, GEMIN4, UNC45A, YME1L1, ARF5, NOP56, EIF2S2, RPL27 |
| K258R | NUP93, JAK1, NCAPD3, GLUD1, CHCHD3, SPATA5, CAND1, TMEM209, PLEKHG4, RPP30, HACD3, ILVBL, SMC4, RPSKA4, CAD, ALDH1B1, RPN1, PPP1CC, ATP1A1, HERC5, RTCB |

Mutant K258R IN bound to the majority of previously reported host factors, but several binding partners were identified as uniquely binding to the K258R mutant IN. Two factors involved in mitotic chromosome condensation (NCAPD3 and SMC4) were found to preferentially bind K258R mutant IN along with multiple components of the catalytic core of the protein phosphatase I (PPI) complex. These factors have clear links to heterochromatin formation and regulation. In addition, gene ontology analysis of the partners revealed an enrichment for genes involved in tRNA processing as well as antiviral interferon stimulated genes (Table 4). It is not immediately obvious how preferential binding of the mutant IN to these proteins would so strongly redirect integrations to centromeric regions.

**Table 4:** Gene ontology analysis of integrase interacting host factors

| GO category | # of proteins | P-value | Proteins |
| --- | --- | --- | --- |
| ***WT (all partners)*** |  |  |  |
| Nucleotide binding | 19 | 5.4E-5 | PRKDC, MDN1, CKMT1A, AIFM1, ABCD3, ATP2A2, ABCE1, SUCLA2, COQ8B, ATAD3C, DDX20, AFG3L2, ATAD3A, ATAD3B, TYK2, MCM7, ARF4, YME1L1, ARF5 |
| ***WT (unique)*** |  |  |  |
| rRNA processing | 4 | 1.7E-3 | TEX10, GEMIN4, NOP56, RPL27 |
| ***K258R (all partners)*** |  |  |  |
| Nucleotide binding | 26 | 5.2E-8 | JAK1, GLUD1, SPATA5, SMC4, RPS6KA4, CAD, ALDH1B1, ATP1A1, RTCB, PRKDC, MDN1, CKMT1A, AIFM1, ABCD3, ATP2A2, ABCE1, SUCLA2, COQ8B, ATAD3C, DDX20, AFG3L2, ATAD3A, ATAD3B, TYK2, MCM7, ARF4 |
| Antiviral mechanism by IFN-stimulated genes | 6 | 2.1E-4 | NUP93, JAK1, HERC5, NUP205, ABCE1, NUP188 |
| tRNA processing in the nucleus | 5 | 8.3E-4 | NUP93, RPP30, RTCB, NUP205, NUP188 |
| PTW/PP1 complex | 2 | 4.9E-2 | PPP1CB, PPP1CC |
| ***K258R (unique)*** |  |  |  |
| tRNA processing | 3 | 2.3E-2 | RPP30, RTCB, NUP93 |
| ISG15 antiviral mechanism | 3 | 4.7E-2 | JAK1, HERC5, NUP93 |
| Meiotic chromosome condensation / condensin complex | 2 | 5.2E-3 | NCAPD3, SMC4 |

To our knowledge, the K258R mutant shows the most dramatic retargeting of integration sites reported for any retrovirus so far. The striking redirection of integrations to the centromere caused by the K258R mutation in the IN protein is especially provocative in light of recent work linking centromeric HIV-1 integrations to viral latency and control. Proviruses in centromeric satellite DNA have been found in the latent reservoir of patients as well as associated with deep viral latency in past reactivation studies^32,34^. Thus, integration into these “gene deserts” promotes viral silencing and the formation of the major impediment to HIV-1 cure. More recently it has been shown that proviral sequences from elite controllers were also preferentially enriched in centromeric satellite DNA^35^, suggesting that a common process may underlie the resultant proviral silencing in both settings. It is not yet known whether IN mutations are associated with increased centromeric integration in patients, but we have found the K258R mutation present at low frequency in proviral sequence repositories of latent proviruses, drug resistant mutants, and from patients on suppressive antiretroviral therapy^36^. Understanding how this single point mutation can cause such a striking retargeting of integration will be important for characterizing and ultimately manipulating the mechanisms that underlie viral latency and long term control in patients.

## Methods

### Cells and plasmids

HEK293T cells and HeLa cells were cultured in DMEM media supplemented with 10% FBS and 1% pen-strep at 37°C, 5% CO_2_.

HIV-1 viral constructs were derived from the replication defective pNL4.3R-E-plasmid (NIH AIDS Reagent Program #3148) carrying a firefly luciferase reporter gene in the *nef* open reading frame. Mutations were introduced into the IN open reading frame using PCR site-directed mutagenesis with custom primer^22^.

### Transfection, virus preparation and infection

To prepare pseudotyped virus for infection, HEK293T cells were co-transfected with the pNL4.3.Luc.R-E-viral vector as well as a plasmid expressing the vesicular stomatitis virus glycoprotein (VSV-G) envelope (pMD2.G) using Lipofectamine 3000 (Life Technologies) according to basic manufacturer’s protocol. Viral supernatants were collected at 48 hours post-transfection, filtered through a 0.45 micron filter, and DNase treated to eliminate plasmid DNA contamination. Viral preparations were normalized by RNA viral genome content, diluted 3-fold with fresh culture medium and immediately used for infection of HeLa cells.

### Luciferase assay

Successful viral transduction was assayed after 48 hours by measuring luciferase activity with the Promega Luciferase Assay System (Cat# E4550). Luminescence (RLU) measurements were normalized for total cell count as determined by protein concentration.

### Quantitative PCR for viral DNA intermediate and RNA analysis

DNA was isolated from acutely infected cells 2 days post-infection using the Qiagen DNeasy Blood and Tissue Kit. Quantitative PCR for viral DNAs was performed using FastStart Universal SYBR Green Mastermix (Bio-Rad) according to manufacturer’s protocol on ABI 7500 Fast Real Time PCR System. Total viral DNA was quantified using primers complementary to the luciferase gene. Reverse transcription (RT) products were detected with LTR specific primers. 2-LTR circles were quantified as previously published and normalized to total virus ^37^. Integrated proviruses were quantified using the published Alu-gag nested PCR protocol ^38,39^.

To quantify steady state viral mRNA levels, RNA was extracted from cells using a standard Trizol protocol. Reverse transcription was performed using random hexamer primers with Maxima H Reverse Transcriptase (Thermo Fisher). Viral cDNA was then quantified via qPCR using primers complementary to spliced *tat* message and normalized to a housekeeping gene.

All primers used for quantification can be found in Table S1. A minimum of three biological replicates were performed per experiment with technical duplicates within each experiment for precision. Biological replicates refer to completely independent experiments, while technical replicates refer to repeated measures of the same samples. A single factor ANOVA analysis was used to identify significant changes (p < 0.05). If appropriate, pairwise comparisons were performed using a two-tailed paired t-test assuming unequal variance.

### Next generation sequencing (NGS) library construction

DNA sequencing libraries were prepared as described previously ^22,25,40^. Briefly, five micrograms of purified genomic DNA from infected cells was randomly sheared using a Branson 450 Digital Sonifier. Sheared ends of DNA were subsequently repaired, A-tailed and ligated to custom oligonucleotide adaptors. Nested PCR was performed using viral and adaptor specific primers to enrich the library for proviral-host genome junctions and add necessary index and flow cell attachment sequences for Illumina (See Table S2 for library adaptor and primer sequences). PCRs were performed such that the final library product should contain 40 bp of the 3′ viral LTR sequence immediately prior to the junction with the host genome sequence. Sequencing was performed using the Illumina MiSeq platform. Three unique biological replicate libraries were generated and sequenced independently.

### Integration site mapping data analysis

Reads were initially demultiplexed by unique dual barcodes and filtered to exclude reads not containing an initial viral LTR sequence at the host junction using a custom python script ^22^. We required an exact match to the terminal 40 nt of the 3′ viral LTR. All reads were then trimmed to remove both leading viral sequence as well as any residual adaptor sequences. Reads of less than 20 nucleotides after all filtering steps were discarded. Remaining reads were mapped to the GRCh38 human genome using either Bowtie2 or BLAT^27,41^.

For majority of analyses, unless otherwise noted, reads were first aligned to the pNL4.3.Luc.R-E-vector genome to remove any viral auto-integration or circular products. The remaining reads were then aligned to the unmasked GRCh38 human reference genome using Bowtie2 end-to-end alignment with a seed length of 28 nucleotides and a maximum of 2 mismatches permitted in the seed. Reads that mapped to multiple locations were not suppressed. Instead, best alignment was reported. For reads with equally good alignments, one of the alignments was reported at random.

Where noted, sequences were further locally aligned to the unmasked GRCh38 genome build using either Bowtie2 sensitive local settings or BLAT. For BLAT analysis, alignments were filtered for 95% minimum identity and a minimum score of 30. All acceptable alignments above this threshold were reported with scores based on number of matched/mismatched bases and a default gap penalty. For reads mapping to multiple locations equally well, all alignments were reported. Parameters for Bowte2 local mapping were 20 nt seed length, allowing 0 mismatches in the seed.

Reads were also aligned directly to the RepeatMasker genome track from UCSC using the same mapping algorithms. Only data from Bowtie2 local mapping is shown here. The RepeatMasker track contains all annotated repeat sequences in the human genome ^31^. Number of integrations falling into each specified repeat class was calculated and presented as a percent of the total number of integrations mapped.

### Hot-spot analysis of viral integrations

Using a previously reported custom perl script, common sites, or “hot-spots”, of viral integration were determined ^42,43^. First, identical reads, or PCR duplicates were condensed. Second, reads with identical junctions but varying sonication breakpoints were condensed to eliminate any confounding effects of clonal expansion. To be stringent, reads with highly similar sequences (i.e. >95% identity) were also combined to eliminate any artifacts produced from small PCR or sequencing errors. From here, “hot-spots” of viral integration were determined using a sliding window approach ^23^. This script searches for multiple integrations falling within a set range of nucleotides from each other. For this study “hot-spots” were defined as regions of 10 kb or less with five or more unique viral integrations.

### Analysis of integration sites with respect to genomic annotations

Genomic coordinates of annotated RefSeq genes, transcription start sites, CpG islands and DNase hypersensitivity regions were extracted from the GRCh38 genome assembly via the UCSC Genome Browser. The genomic coordinates of centromeric sequences were also extracted from UCSC Genome Browser. Locations of RNA polymerase II binding sites and histone modifications were extracted from ENCODE data sets generated from uninfected HeLa cells (Pol II: ENCFF246QVY; H3K27Ac: ENCFF113QJM ; H3K9me3: ENCFF712ATO ; H3K36me3: ENCFF864ZXP ; H3K4me3: ENCFF862LUQ). Distance of proviral integrations to nearest feature was calculated using BedTools 44. A matched random control (MRC) data set of comparable size was generated with BedTools Random command and mapped in parallel to experimental data sets.

A one-sample t-test was used to compare integration distribution between experimental samples and MRC (Table S3). To gauge the statistical significance of differences in integration patterns between WT IN and mutant IN we used a paired t-test of three independent replicate data sets for each condition or Fisher’s exact test on the aggregate integration data (Table S4).

### Sequence analysis of centromeric integration sites

The host sequence flanking the site of integration was extracted from Bed coordinates of mapped integration sites. To align sites of integration along the repeat length of the alphoid repeat, we used only the 5 base pairs flanking the site of integration (total length 10 bp) to align to a consensus sequence for the alphoid repeat monomer (AJ131208.1). Only unique junctions were aligned. Alignments were performed with Clustal Omega ^45^. For count purposes, we defined 17 bins spanning the alphoid repeat monomer, each consisting of ten base pairs, and counted the number of integrations falling in each bin.

### PCR assays for quantifying centromeric integrations

To determine if centromeric DNA sequences were over-represented in library preparations, we made use of previously reported unique chromosome specific centromere primers ^46^. Amplified viral-host genome fragments from library preparations were used in a qPCR assay using centromere specific primers to relatively compare quantities of centromeric DNA sequences between infections with viruses carrying WT or mutant IN proteins.

To look more generically at integration into all centromeres, we devised a nested PCR assay based on both the basic Alu-gag PCR protocol for quantifying proviral integration and a previously published assay using alpha satellite specific primers (alphoid-1, alphoid-2) ^26^. For the first nest, one of two primers complementary to the alpha satellite consensus sequence were used in conjunction with either a 5′ viral specific primer (5′-gag) or a 3′ viral specific primer (3′-luc). For validation purposes, a number of randomly selected fragments were cloned from the first rounds of PCR and sequenced by Sanger sequencing to verify that we were indeed amplifying alphoid repeats at the viral-host genome junction. LTR-specific primers were then used for the second nest quantitative PCR. These values were normalized to total LTR content in original unamplified DNA. See Table S1 for primer sequences used.

### Co-immunoprecipitation of IN proteins and mass spectrometry

Either WT HIV-1 IN or IN harboring the K258R mutation was cloned into a mammalian expression vector (pJET). Both proteins had an N-terminal HA-tag for immunoprecipitation. As a negative control, we also transfected cells with an empty HA-vector. Constructs were transfected into HEK293T cells as described. After 24 hours, cells were collected, washed and lysed with an NP-40 lysis buffer (20 mM Tris HCl, pH 8; 137 mM NaCl, 2 mM EDTA and 1% NP-40). Adequate, comparable expression of WT and mutant IN proteins was confirmed via Western blot using HA-specific or IN-specific antibodies.

Cell lysates were subsequently mixed with BSA blocked HA-coated magnetic beads (Pierce) and rotated overnight at 4°C. Beads were washed three times with lysis buffer, finished with two PBS washes and sent for mass spectrometry analysis (Rockefeller Mass Spectrometry Core Facility).

MS results were filtered by number of peptides detected vs. an empty HA vector control. Only proteins with five or more spectral counts were considered. Proteins were considered enriched when there was a minimum of 5-fold more unique spectral counts detected in the IN immunoprecipitation vs. the control precipitation. Enriched peptides immunoprecipitated by WT and K258R mutant IN were further subjected to gene ontology analysis performed with gProfiler software ^47^.

### Data and code availability

Sequencing reads generated as part of this study are available at the NCBI Sequencing Read Archive: XXXX. Code uniquely generated for this analysis is available upon request.

## Acknowledgements

This study was supported by NCI grant R01 CA 30488 from the National Cancer Institute (S.P.G) and NIAID Ruth L. Kirschstein NRSA fellowship F32 AI 149989 from the National Institute of Allergy and Infectious Disease (S.W.). S.P.G. is an Investigator of the Howard Hughes Medical Institute.

## Author contributions

Conceptualization and methodology: SW and SPG; Data curation, formal analysis and visualization: SW; Supervision, project administration and resources: SPG; Funding acquisition: SW and SPG; Writing – original draft: SW; Writing - review and editing: SPG and SW

## Additional Information

Supplementary Information is available for this paper.

Correspondence and requests for materials should be addressed to Stephen P. Goff.

## Ethics declarations

The authors declare no competing interests.

## Notes

### Competing Interest Statement

The authors have declared no competing interest.

## References

1. Bowerman, B., Brown, P. O., Bishop, J. M. & Varmus, H. E. A nucleoprotein complex mediates the integration of retroviral DNA. Genes Dev. 3, 469–478 (1989).

2. Brown, P. O., Bowerman, B., Varmus, H. E. & Bishop, J. M. Retroviral integration: Structure of the initial covalent product and its precursor, and a role for the viral IN protein. Proc. Natl. Acad. Sci. U. S. A. 86, 2525–2529 (1989).

3. Bor, Y. C., Bushman, F. D. & Orgel, L. E. In vitro integration of human immunodeficiency virus type 1 cDNA into targets containing protein-induced bends. Proc. Natl. Acad. Sci. U. S. A. 92, 10334–8 (1995).

4. Craigie, R., Fujiwara, T. & Bushman, F. The IN protein of Moloney murine leukemia virus processes the viral DNA ends and accomplishes their integration in vitro. Cell 62, 829–837 (1990).

5. Brown, P. O., Bowerman, B., Varmus, H. E. & Bishop, J. M. Correct integration of retroviral DNA in vitro. Cell 49, 347–356 (1987).

6. Mitchell, R. S. et al.. Retroviral DNA integration: ASLV, HIV, and MLV show distinct target site preferences. PLoS Biol. 2, E234 (2004).

7. Schröder, A. R. W. et al.. HIV-1 integration in the human genome favors active genes and local hotspots. Cell 110, 521–9 (2002).

8. Debyser, Z., Christ, F., De Rijck, J. & Gijsbers, R. Host factors for retroviral integration site selection. Trends Biochem. Sci. 40, 108–16 (2015).

9. Ciuffi, A. et al.. A role for LEDGF/p75 in targeting HIV DNA integration. Nat. Med. 11, 1287–9 (2005).

10. Llano, M. et al.. An essential role for LEDGF/p75 in HIV integration. Science (80-.). 314, 461–4 (2006).

11. Maertens, G. et al.. LEDGF/p75 is essential for nuclear and chromosomal targeting of HIV-1 integrase in human cells. J. Biol. Chem. 278, 33528–33539 (2003).

12. McNeely, M. et al.. In vitro DNA tethering of HIV-1 integrase by the transcriptional coactivator LEDGF/p75. J. Mol. Biol. 410, 811–830 (2011).

13. Kvaratskhelia, M., Sharma, A., Larue, R. C., Serrao, E. & Engelman, A. Molecular mechanisms of retroviral integration site selection. Nucleic Acids Res. 42, 10209–25 (2014).

14. Lesage, P. & Todeschini, A. L. Happy together: The life and times of Ty retrotransposons and their hosts. Cytogenetic and Genome Research vol. 110 70–90 (2005).

15. Zou, S., Ke, N., Kim, J. M. & Voytas, D. F. The Saccharomyces retrotransposon Ty5 integrates preferentially into regions of silent chromatin at the telomeres and mating loci. Genes Dev. 10, 634–645 (1996).

16. Xie, W. et al.. Targeting of the Yeast Ty5 Retrotransposon to Silent Chromatin Is Mediated by Interactions between Integrase and Sir4p. Mol. Cell. Biol. 21, 6606–6614 (2001).

17. Dai, J., Xie, W., Brady, T. L., Gao, J. & Voytas, D. F. Phosphorylation Regulates Integration of the Yeast Ty5 Retrotransposon into Heterochromatin. Mol. Cell 27, 289–299 (2007).

18. Andrake, M. D. & Skalka, A. M. Retroviral Integrase: Then and Now. Annu. Rev. Virol. 2, 241–264 (2015).

19. Chen, L., Keppler, O. T. & Schölz, C. Post-translational modification-based regulation of HIV replication. Frontiers in Microbiology vol. 9 (2018).

20. Cereseto, A. et al.. Acetylation of HIV-1 integrase by p300 regulates viral integration. EMBO J. 24, 3070–3081 (2005).

21. Terreni, M. et al.. GCN5-dependent acetylation of HIV-1 integrase enhances viral integration. Retrovirology 7, (2010).

22. Winans, S. & Goff, S. P. Mutations altering acetylated residues in the CTD of HIV-1 integrase cause defects in proviral transcription at early times after integration of viral DNA. PLOS Pathog. 16, e1009147 (2020).

23. Berry, C. C., Ocwieja, K. E., Malani, N. & Bushman, F. D. Comparing DNA integration site clusters with scan statistics. Bioinformatics 30, 1493–1500 (2014).

24. Justice, J. et al.. The MET Gene Is a Common Integration Target in Avian Leukosis Virus Subgroup J-Induced Chicken Hemangiomas. J. Virol. 89, 4712–4719 (2015).

25. Malhotra, S. et al.. Selection for avian leukosis virus integration sites determines the clonal progression of B-cell lymphomas. PLoS Pathog 13, (2017).

26. Carteau, S., Hoffmann, C. & Bushman, F. Chromosome Structure and Human Immunodeficiency Virus Type 1 cDNA Integration: Centromeric Alphoid Repeats Are a Disfavored Target. J. Virol. 72, 4005–4014 (1998).

27. Kent, W. J. BLAT---The BLAST-Like Alignment Tool. Genome Res. 12, 656–664 (2002).

28. McNulty, S. M. & Sullivan, B. A. Alpha satellite DNA biology: finding function in the recesses of the genome. Chromosome Research vol. 26 115–138 (2018).

29. Miga, K. H. Centromeric satellite DNAs: Hidden sequence variation in the human population. Genes vol. 10 (2019).

30. Hartley, G. & O’neill, R. J. Centromere repeats: Hidden gems of the genome. Genes vol. 10 (2019).

31. Tarailo-Graovac, M. & Chen, N. Using RepeatMasker to identify repetitive elements in genomic sequences. Current Protocols in Bioinformatics (2009) doi:10.1002/0471250953.bi0410s25.

32. Jordan, A., Bisgrove, D. & Verdin, E. HIV reproducibly establishes a latent infection after acute infection of T cells in vitro. EMBO J. 22, 1868–1877 (2003).

33. Jäger, S. et al.. Global landscape of HIV-human protein complexes. Nature 481, 365–370 (2012).

34. Lewinski, M. K. et al.. Retroviral DNA integration: viral and cellular determinants of target-site selection. PLoS Pathog. 2, e60 (2006).

35. Jiang, C. et al.. Distinct viral reservoirs in individuals with spontaneous control of HIV-1. Nature (2020) doi:10.1038/s41586-020-2651-8.

36. Foley, B. et al.. HIV Sequence Compendium 2018. in (Theoretical Biology and Biophysics Group, Los Alamos National Laboratory, NM, LA-UR 18-25673).

37. Mandal, D. & Prasad, V. R. Analysis of 2-LTR circle junctions of viral DNA in infected cells. Methods Mol. Biol. 485, 73–85 (2009).

38. Butler, S. L., Hansen, M. S. & Bushman, F. D. A quantitative assay for HIV DNA integration in vivo. Nat. Med. 7, 631–4 (2001).

39. O’Doherty, U., Swiggard, W. J., Jeyakumar, D., McGain, D. & Malim, M. H. A Sensitive, Quantitative Assay for Human Immunodeficiency Virus Type 1 Integration. J. Virol. 76, 10942–10950 (2002).

40. Serrao, E., Cherepanov, P. & Engelman, A. N. Amplification, next-generation sequencing, and genomic DNA mapping of retroviral integration sites. J. Vis. Exp. 2016, (2016).

41. Langmead, B., Trapnell, C., Pop, M. & Salzberg, S. L. Ultrafast and memory-efficient alignment of short DNA sequences to the human genome. Genome Biol. 10, R25 (2009).

42. Justice, J. F., Morgan, R. W. & Beemon, K. L. Common Viral Integration Sites Identified in Avian Leukosis Virus-Induced B-Cell Lymphomas. MBio 6, e01863–15 (2015).

43. Malhotra, S. et al.. Selection for avian leukosis virus integration sites determines the clonal progression of B-cell lymphomas. PLOS Pathog. 13, e1006708 (2017).

44. Quinlan, A. R. & Hall, I. M. BEDTools: A flexible suite of utilities for comparing genomic features. Bioinformatics 26, 841–842 (2010).

45. Sievers, F. & Higgins, D. G. Clustal omega, accurate alignment of very large numbers of sequences. Methods Mol. Biol. 1079, 105–116 (2014).

46. Contreras-Galindo, R. et al.. Rapid molecular assays to study human centromere genomics. Genome Res. 27, 2040–2049 (2017).

47. Reimand, J., Kull, M., Peterson, H., Hansen, J. & Vilo, J. g:Profiler--a web-based toolset for functional profiling of gene lists from large-scale experiments. Nucleic Acids Res. 35, W193–200 (2007).

