## Supplementary material for "A point mutation in HIV-1 integrase redirects proviral integration into centromeric repeats": Fig S1

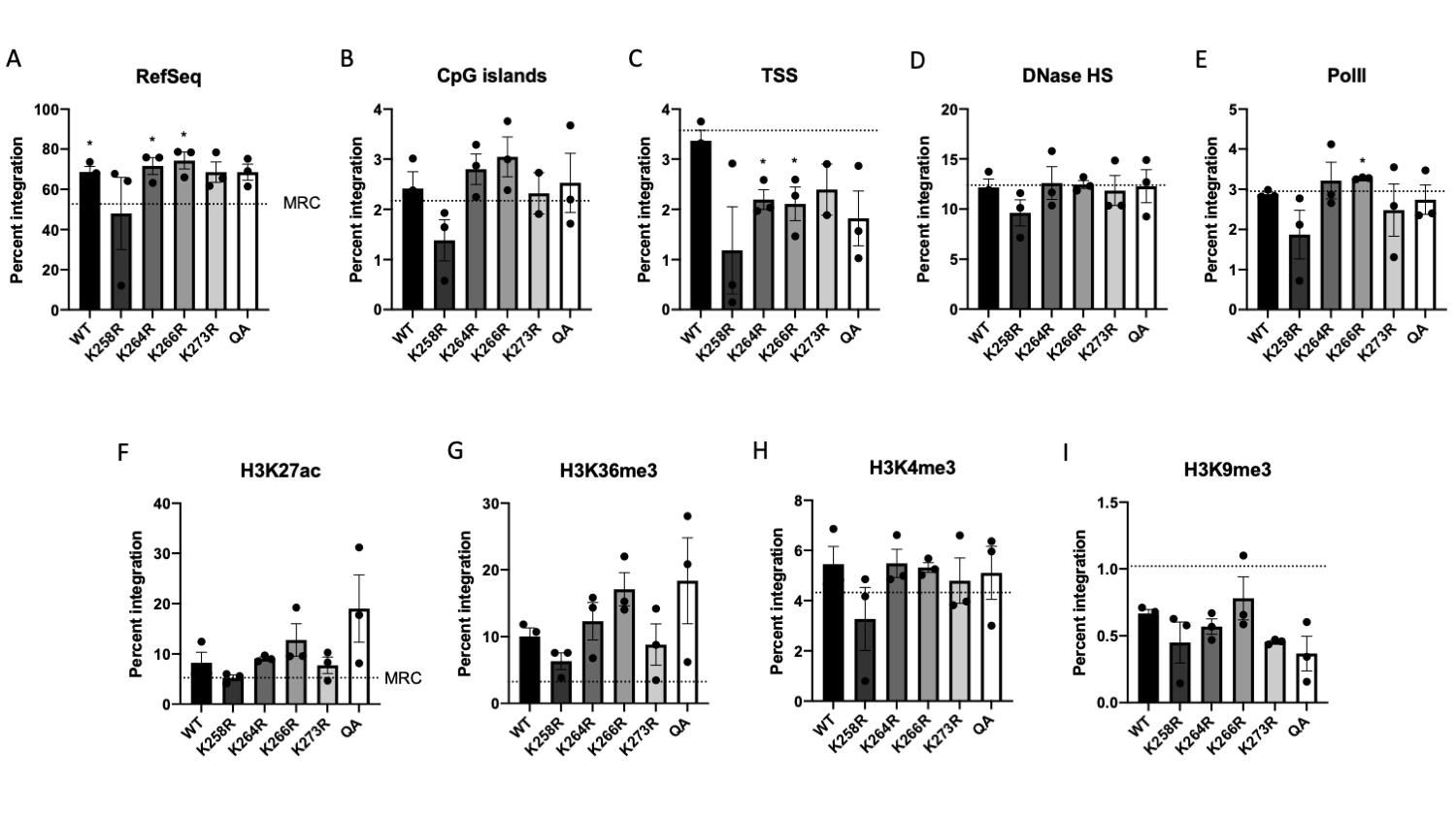

**Figure S1: Integration frequency of WT and acetylation mutant IN proteins with respect to common genomic features**. Frequency of integrations falling within 1 kb of (A) RefSeq genes, (B) CpG islands, (C) transcription start sites, (D) DNase hypersensitivity sites, (E) RNA polymerase II binding sites and various pre-infection histone modification sites in HeLa cells (F-I) was calculated using BedTools. Frequency of integrations in a matched random control (MRC) data set is shown as dashed line. Data is shown as the average of three independent replicates +/- SEs. Statistical significance of integration frequency relative to MRC was gauged by a one-sample t-test (all p-values shown in Table S3).
