## Supplementary material for "A point mutation in HIV-1 integrase redirects proviral integration into centromeric repeats": Fig S2

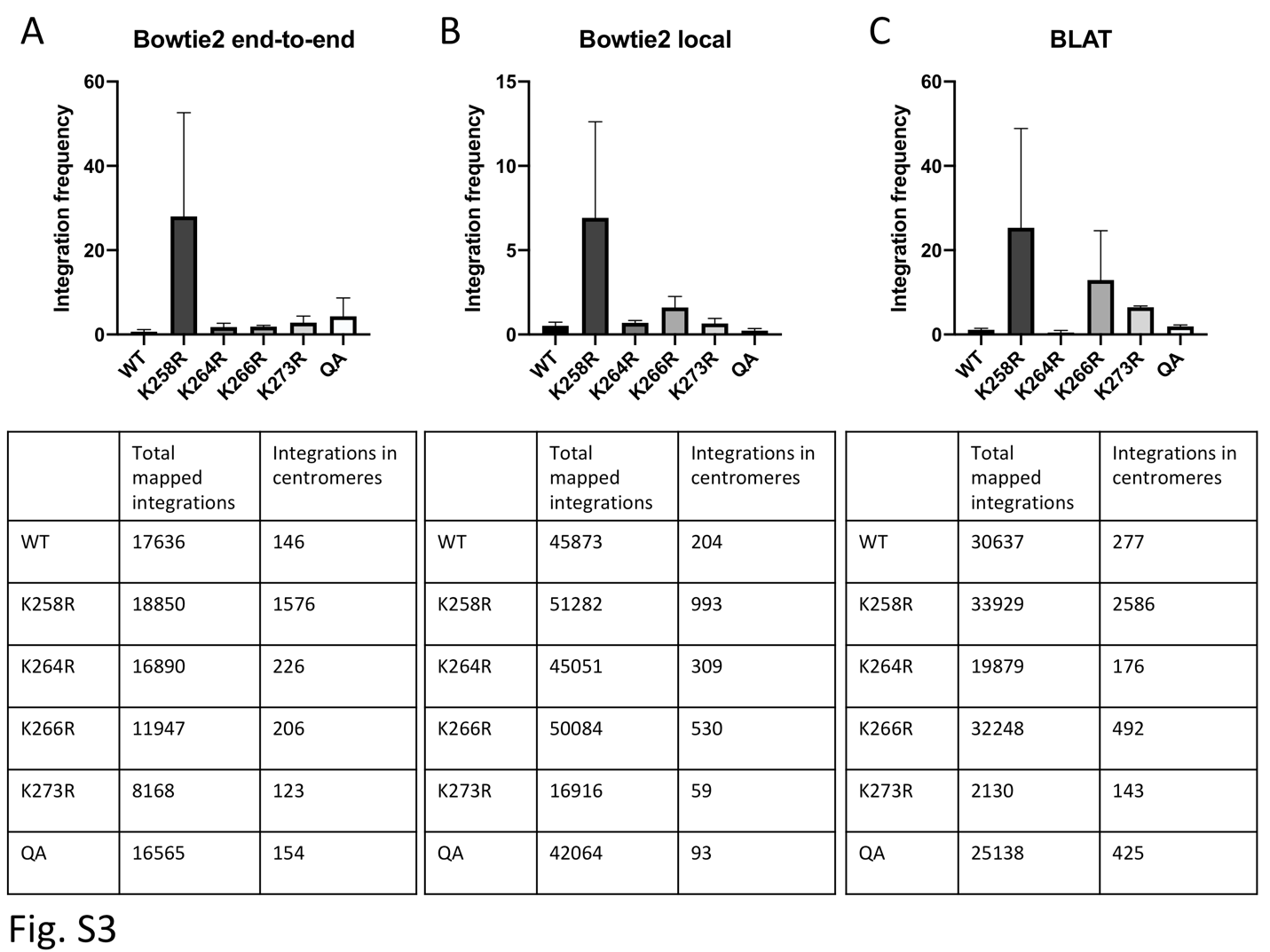

**Figure S2: Integration frequency into centromeres using different mapping algorithms**. NGS data from three independent biological replicates was mapped to the GRCh38 human genome assembly using (A) Bowtie2 end-to-end alignment (shown in main text), (B) Bowtie2 sensitive local alignment or (C) BLAT alignment algorithms. Bar graphs show the integration frequency into centromeres as determined by each algorithm. Graphed data is presented as an average of three independent biological replicates +/- SEs. Absolute number of detected unique integrations for all libraries summed is shown below each graph.
