## Supplementary material for "A point mutation in HIV-1 integrase redirects proviral integration into centromeric repeats": Table S1

**Table S1**: Primer sequences used for quantitative PCR analysis of viral DNA intermediates and transcripts.

| **Target** | **Primer sequence (5’-3’)** |
| --- | --- |
| Late RT | TGTGTGCCCGTCTGTTGTGT |
|  | GAGTCCTGCGTCGAGAGATC |
| Luciferase | CGTCTTTCCGTGCTCCAAAAC |
|  | CAAAGGATATCAGGTGGCCC |
| 2LTR circles | AACTAGGGAACCCACTGCTTAAG |
|  | TCCACAGATCAAGGATATCTTGTC |
| Alu-gag nest 1 | GCCTCCCAAAGTGCTGGGATTACAG |
|  | GCTCTCGCACCCATCTCTCTCC |
| Alu-gag nest 2 | GCCTCAATAAAGCTTGCCTTGA |
|  | TCCACACTGACTAAAAGGGTCTGA |
| *Tat* mRNA | GTTTGTTTCATGACAAAAGCCTTA |
|  | CTATTCCTTCGGGCCTGTC |
| Chr1 | GTTCCCTTAGACAGAGCAGATTT |
|  | CAACGCAGTTTGTGGGAATG |
| Chr2 | TCGTTGGAAACGGGATTGT |
|  | CTGCTCTATGAAAGGGACTGTT |
| Chr4 | CTGTAGTATCTGGAAGTGGACATT |
|  | GGTTCAACTGTGTTCGTTTAGG |
| Chr14 | GATTTCGTTGGAAACGGGATTAC |
|  | AGAAAGATCCACGCCTGTTA |
| Alphoid-1 | GCAAGGGGATATGTGGACC |
| Alphoid-2 | ACCACCGTAGGCCTGAAAGCAGTC |
| 5’-gag | GCTCTCGCACCCATCTCTCTCC |
| 3’-luc | AGGCCAAGAAGGGCGGAAAG |
