## Supplementary material for "A point mutation in HIV-1 integrase redirects proviral integration into centromeric repeats": Table S2

**Table S2**: Adaptor and primer sequences used for construction of integration site mapping NGS libraries.

| **Primer name** | **Primer sequence** |
| --- | --- |
| Adaptor short arm | P-GATCGGAAGAGCAAAAAAAAAAAAAAAA |
| Adaptor long arm | CAAGCAGAAGACGGCATACGAGATnnnnnnGTGACTGGAGTTCAGACGTGTGCTCTTCCGATC*T |
| PCR-1-F | TGTGACTCTGGTAACTAGAGATCCCTC |
| PCR-1-R | CAAGCAGAAGACGGCATACGAGAT |
| PCR-2-F | AATGATACGGCGACCACCGAGATCTACACTCTTTCCCTACACGACGCTCTTCCGATCTGAGATCCCTCAGACCCTTTTAGTCAG |
| PCR-2-R | CAAGCAGAAGACGGCATACGAGATnnnnnn |

nnnnnn denotes a 6-bp unique barcode, P denotes phosphorylation and * denotes a phosphorothioate bond
