## Supplementary material for "A point mutation in HIV-1 integrase redirects proviral integration into centromeric repeats": Table S3

**Table S3**: Statistical analysis of integration preferences of WT or mutant IN proteins as compared to a matched random control (one sample t-test, 1 kb window).

|  | **WT** | **K258R** | **K264R** | **K266R** | **K273R** | **QA** |
| --- | --- | --- | --- | --- | --- | --- |
| RefSeq genes | 0.0285 | 0.8183 | 0.0463 | 0.0356 | 0.0885 | 0.058 |
| TSS | 0.4319 | 0.1111 | 0.0197 | 0.0489 | 0.2596 | 0.0849 |
| CpG islands | 0.5413 | 0.1959 | 0.1735 | 0.1587 | 0.7776 | 0.6058 |
| RNA Pol II | 0.3183 | 0.2166 | 0.6193 | 0.0018 | 0.5446 | 0.6232 |
| DNase HS | 0.8181 | 0.1693 | 0.9057 | 0.8804 | 0.7553 | 0.6058 |
| H3K27ac | 0.2927 | 0.9830 | 0.0079 | 0.1451 | 0.2675 | 0.1759 |
| H3K36me3 | 0.0336 | 0.1331 | 0.0837 | 0.0304 | 0.2136 | 0.1432 |
| H3K4me3 | 0.2517 | 0.4915 | 0.1764 | 0.0356 | 0.6527 | 0.535 |
| H3K9me3 | 0.0073 | 0.0653 | 0.016 | 0.2742 | 0.0003 | 0.0372 |
| Centromeric | 0.1580 | 0.3991 | 0.9515 | 0.8294 | 0.598 | 0.6204 |
