## Supplementary material for "A point mutation in HIV-1 integrase redirects proviral integration into centromeric repeats": Table S4

**Table S4**: Statistical analysis of integration preferences of K258R mutant IN protein as compared to WT (paired t-test or Fisher’s exact test, 1 kb window).

|  | **Paired t-test (N=3)** | **Fisher’s exact test** |
| --- | --- | --- |
| RefSeq genes | 0.3247 | <0.0001 |
| TSS | 0.1691 | <0.0001 |
| CpG islands | 0.1547 | <0.0001 |
| RNA Pol II | 0.2656 | 0.0051 |
| DNase HS | 0.3578 | 0.0.006 |
| H3K27ac | 0.3797 | <0.0001 |
| H3K36me3 | 0.2531 | <0.0001 |
| H3K4me3 | 0.3805 | 0.0025 |
| H3K9me3 | 0.2278 | 0.1847 |
| Centromeric | 0.3860 | <0.0001 |
